# Real-time environmental monitoring of contaminants using living electronic sensors

**DOI:** 10.1101/2021.06.04.447163

**Authors:** Joshua T. Atkinson, Lin Su, Xu Zhang, George N. Bennett, Jonathan J. Silberg, Caroline M. Ajo-Franklin

## Abstract

Real-time chemical sensing is needed to counter the global threats posed by pollution. We combine synthetic biology and materials engineering to develop a living bioelectronic sensor platform with minute detection times. *Escherichia coli* was programmed to reduce an electrode in a chemical-dependent manner using a modular, eight-component, synthetic electron transport chain. This strain produced significantly more current upon exposure to thiosulfate, an anion that causes microbial blooms. Incorporating a protein switch into the synthetic pathway and encapsulation of microbes with electrodes and conductive nanomaterials yielded a living bioelectronic sensor that could detect an endocrine disruptor within two minutes in riverine water, implicating the signal as mass transfer limited. These findings provide a new platform for miniature, low-power sensors that safeguard ecological and human health.

**One Sentence Summary:** Chemicals are detected electrically using an allosterically-regulated electron transfer pathway in designer microbes.

## MAIN TEXT

Contamination of freshwater with natural and synthetic chemicals is a global environmental challenge (*1*). Of particular concern are chemicals that affect vertebrate reproduction and inorganic compounds that stimulate microbial blooms, as both can have severe ecological impacts when they enter the environment (*2*–*4*). Because chemical releases can be dynamic and transient, there is a need to sense these chemicals *in situ*, in real time (*4*). This sensing must also be accurate across environments with varying abiotic conditions.

Recent strides in biosensing enable the detection of contaminants via different modalities. Synthetic biology has yielded field-deployable biosensors that monitor chemical contaminants (*5*), reporting them as visual signals (*6*, *7*). Alternatively, bioelectronic sensors have been developed using bacteria that couple chemical sensing to the production of electrical current through a process called extracellular electron transfer (EET) (*8*–*12*). These sensors all rely on regulating transcription for detection, limiting their response times to ≥30 minutes.

Engineered microorganisms have been integrated into materials to create free-standing devices for sensing diverse chemicals (*13*). These approaches, which typically encapsulate bacteria in hydrogels, have yielded deployable optical sensors for explosives (*14*), heavy metals (*15*), and chemical inducers (*16*, *17*). While providing mechanical integrity and supporting continuous sensing, these materials attenuate transmission of the signals, degrading the signal-to-noise ratio and temporal response.

Thus, to enable real-time environmental biosensing of chemicals, we need new strategies to rapidly control and robustly transmit electrical current from microbes to electronics. Here, we combine synthetic biology and materials engineering to overcome these challenges in parallel by programming a microbe to report in real time on contaminants that trigger rapid microbial growth and impair vertebrate reproduction, interfacing these cells with electrodes using synthetic materials to enhance the signal-to-noise of conditional EET, and showing that this bioelectronic sensor platform can sense different chemicals in urban waterway samples.

To develop a strategy to rapidly report on inorganic nutrients that trigger microbial blooms by producing current, we designed a synthetic electron transfer (ET) pathway in *Escherichia coli* where sulfur oxyanions gate electron flow to an electrode. We chose to test this strategy using thiosulfate, a common dechlorination agent used in water treatment that can trigger microbial blooms that impact aquatic habitats when used in excess (*2*). We designed a thiosulfate-dependent ET pathway using three modules (Fig. 1A). The *<u>Input (I)</u>* module couples NADPH oxidation to the reduction of sulfite, a product of sulfur assimilation when cells are exposed to thiosulfate, through the expression of ferredoxin-NADP reductase (FNR), ferredoxin (Fd), and sulfite reductase (SIR). The *<u>Coupling (C)</u>* module uses the product of the Input module, sulfide, and a sulfide-quinone reductase (SQR) to reduce inner-membrane quinones to quinols. Lastly, the <u>*Output (O)*</u> module, composed of the quinol dehydrogenase CymA and the cytochrome *c*-porin complex MtrCAB (CymA-MtrCAB), rapidly transfers electrons from quinols to an electrode (*18*, *19*). These modules route electrons from NADPH to an electrode using an ET pathway that requires twenty-nine cofactors, including twenty-four hemes, two flavins, one siroheme, one 4Fe4S, and one 2Fe2S.

**Fig. 1.**
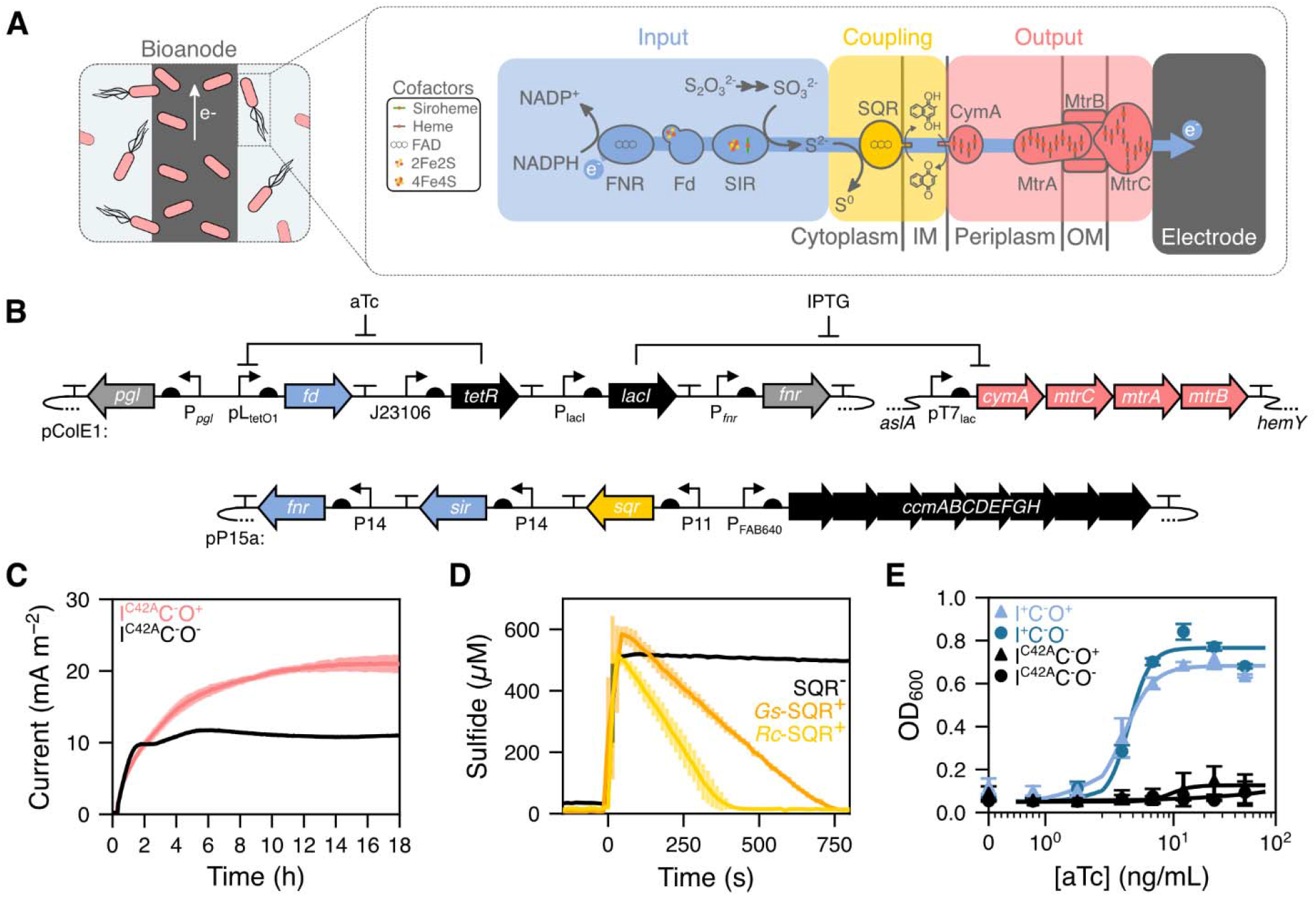
An *E. coli* sensor with a synthetic electron transfer chain. **(A)** Fd-dependent ET from FNR to SIR couples NADPH oxidation to sulfite reduction (Input module), SQR uses sulfide oxidation to reduce quinones (Coupling module), and CymA-MtrCAB use quinol oxidation to drive EET (Output module). **(B)** Genetic circuits for expressing the different modules. **(C)** Current production in a BES containing either I^C42A^C^−^O^+^ (red) or I^C42A^C^−^O^−^ (black). Current from I^C42A^C^−^O^+^ was significantly greater than I^C42A^C^−^O^−^ 3 h after introduction to the BES (*p =* 0.05). **(D)** Sulfide oxidation by I^C42A^C^−^O^−^ cells expressing *Rc*-SQR (yellow, 370 μmol s^−1^) and *Gs-*SQR (orange, 230 μmol s^−1^) are significantly faster than cells lacking SQR (black, 5.7 μmol s^−1^) (*p <* 0.01). **(E)** The optical density of I^+^C^−^O^+^ (blue triangles) I^+^C^−^O^−^ (blue circles) cultures increase with aTc concentration. In contrast, I^+^C^−^O^+^ and I^+^C^−^O^−^ both present significantly higher growth complementation than I^C42A^C^−^O^−^ and I^C42A^C^−^O^+^ (black circles and triangles, respectively) upon Fd induction at ≥3.125 ng/mL aTc (*p <* 0.05). Data represents the mean values with error bars representing one standard deviation (n = 3 biologically independent samples). P values were obtained using two-tailed, independent t-tests.

To evaluate the performance of individual modules, we used a combination of genomic- and plasmid-encoded genetic circuits that allowed for plug-and-play expression of module components (Fig. 1B). The Output module (O^+^) was created by integrating an isopropyl β-d-1-thiogalactopyranoside (IPTG)-inducible operon encoding CymA-MtrCAB from *Shewanella oneidensis* MR-1 into the *E. coli* genome and introducing a plasmid that constitutively expresses the cytochrome *c* maturation (*ccm*) operon (*20*). Strains that express the Input and Coupling modules (I^+^ and C^+^) components were created by introducing modified *ccm* plasmids that constitutively express a subset of the Input module components (FNR and SIR) and the Coupling module (SQR). Additionally, a second plasmid was introduced that expresses the Fd using an anhydrotetracycline (aTc) inducible promoter. An ET-deficient version of the Input module (I^C42A^) was created as a negative control by generating a Fd Cys42Ala mutant that cannot coordinate an iron-sulfur cluster (*21*). To minimize off-pathway ET that competes with the desired electron flux, we used a redox-insulated strain, *E. coli* EW11 (*22*), as our parental strain. Using different combinations of these plasmids (Table S1), a set of strains (Table S2) were constructed to evaluate the activity of the individual modules and their combinations.

To optimize the function of the Output module, we assayed its expression, EET, and effect on cell fitness under varying induction conditions. The production of the Mtr cytochromes (Fig. S1A) and EET (Fig. S1B) peaked between 2 and 12.5 μM IPTG. Greater than 10 μM IPTG decreased I^C42A^C^−^O^+^ growth significantly relative to I^C42A^C^−^O^−^ (Fig. S1C). Thus, for all subsequent studies, we induced cells with 10 μM IPTG to maximize EET while minimizing fitness burdens. Using this optimal induction, we compared I^C42A^C^−^O^+^ and I^C42A^C^−^O^−^ anode reduction in a bioelectrochemical system (BES) containing M9 medium and glucose (Fig. 1C). Within 3 h, the I^C42A^C^−^O^+^ strain produced significantly more current than the I^C42A^C^−^O^−^ strain (p-value = 0.05), demonstrating the optimized Output module is functional.

Our next step was to identify an SQR for the Coupling module that would rapidly oxidize sulfide at the concentrations produced by the Input module. The cellular activities of two SQR homologs were compared using I^C42A^C^+ −^O^−^ strains, including the SQRs from *Rhodobacter capsulatus (Rc*-SQR) and *Geobacillus stearothermophilus* (*Gs-*SQR) (*23*, *24*). When cells expressing *Rc-*SQR or *Gs*-SQR were exposed to exogenous sulfide, oxidation of 500 μM sulfide required 7 and 12.5 min, respectively (Fig. 1D). In contrast, cells lacking an SQR did not consume sulfide. When cells were labeled with a fluorescent probe for intracellular sulfane sulfur, an indication of SQR activity (*25*), both SQR-expressing strains accumulated significantly more sulfane sulfur than the empty vector control (Fig. S2). Because *Rc*-SQR oxidized sulfide at faster rates, it was used as the Coupling module for all subsequent studies.

To investigate whether the Input module acquires iron cofactors required for ET in the presence of the Output module, which has a high iron cofactor demand, we measured ET mediated by the Input module in the I^+^C^−^O^+^ and I^+^C^−^O^−^ strains. We leveraged a prior demonstration that growth of our parental strain can be coupled to sulfite reduction by expressing *Mastigocladus laminosus* Fd and *Zea mays* FNR and SIR (*21*). With this cellular assay, Fd-mediated ET from FNR to SIR is required to synthesize cysteine when sulfate or thiosulfate are provided as a sulfur source (*21*) (Fig. S3A). Fd complementation was similar in I^+^C^−^O^+^ and I^+^C^−^O^−^ cells (Fig. 1E). This finding indicates that cells can synthesize holoproteins in the Input module while expressing the Output module.

We predicted that the minimum thiosulfate concentration that our system could ultimately detect would have to be greater than the thiosulfate needed to meet assimilation needs. To establish this lower limit, we evaluated the effect of varying thiosulfate concentrations on the growth (Fig. S3D) and H_2_S evolution (Fig. S3E) from I^+^C^−^O^−^ cultures. Cells added to medium containing ≤0.25 mM thiosulfate as a sulfur source displayed growth complementation that was dependent on Fd expression. At higher thiosulfate concentrations, cells grew to similar densities, suggesting excess thiosulfate was available. In addition, H_2_S was only observed when >0.5 mM thiosulfate was added. Taken together, these results suggest that ET through our full synthetic pathway should be measurable when thiosulfate is >0.25 mM, which is lower than its EC_50_ for fish (4 mM).

To determine if ET through the full synthetic pathway is dependent upon thiosulfate, we integrated all three modules together to build an I^+^C^+^O^+^ strain and measured thiosulfate-dependent EET of these planktonic cells in a BES. Thiosulfate increased the current response of the I^+^C^+^O^+^ strain relative to the I^C42A^C^+^O^+^ strain (Fig. S4), indicating the full pathway acts as a thiosulfate sensor. However, the signal-to-noise was low, making it challenging to discern the signals from these strains. To investigate if this noise arose from using planktonic cells bound loosely to the anode, we encapsulated each strain with the carbon felt working electrode within an alginate-agarose hydrogel (Fig. 2A–B). Compared to planktonic cells, the encapsulated cells responded to thiosulfate with higher signal-to-noise (>5-fold increase in signal intensity), increased reproducibility (>50% decrease in standard deviation), and enhanced linearity (>10-fold increase in R^2^) (Fig. 2C–D). This encapsulation approach produces a living electronic sensor, which was used in all subsequent experiments.

**Fig. 2.**
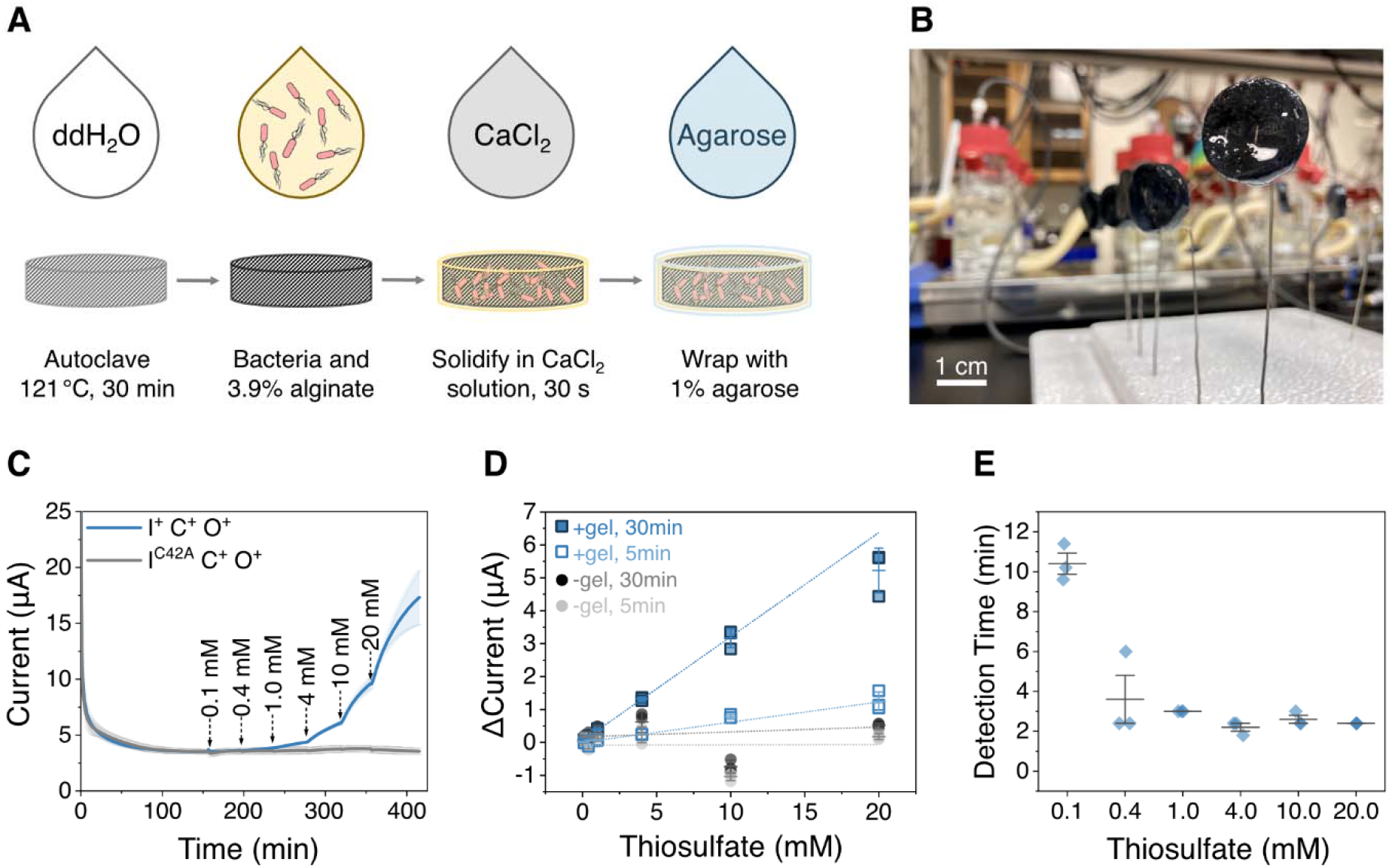
Encapsulation of a living electronic sensor yields rapid detection and quantification of thiosulfate. **(A)** Schematic depicting the protocol for encapsulating the living electronic sensor with an electrode and **(B)** an image of the encapsulated sensor. **(C)** Current generated by the I^+^C^+^O^+^ and I^C42A^C^+^O^+^ strains upon exposure to increasing concentrations of thiosulfate. **(D)** Average current response are a linear function of thiosulfate concentration at 5 (R^2^ = 0.984) and 30 min (R^2^ = 0.994) after thiosulfate addition. **(E)** Detection time for different thiosulfate concentrations (*p <* 0.05). Data represents the mean values with error bars representing one standard deviation (n = 3 biologically independent samples). P values were calculated using a one-way ANOVA with Tukey test.

We next probed the ability of this living electronic sensor to sense different thiosulfate concentrations. Upon addition of as little as 0.1 mM thiosulfate, the I^+^C^+^O^+^ strain immediately presented increased current, while the sensors with the I^C42A^C^+^O^+^ strain did not respond to 20 mM thiosulfate (Fig. 2C). The thiosulfate-dependent signal of the I^+^C^+^O^+^ strain was calculated as the difference between current immediately before each injection (I_t_0_) and current at a fixed time after injection (I_t_injection_). The current response (I_t_injection_ - I_t_0_) is linearly related to the thiosulfate concentration (Fig. 2D) with R^2^ of 0.984 and 0.994 for 5 and 30 min, respectively. By comparing the current differences between the I^C42A^C^+^O^+^ and I^+^C^+^O^+^ strains, thiosulfate was detected with ≥95% confidence within 2 to 10 min of exposure (Fig. 2E). This analysis detects ≥0.4 mM thiosulfate in <4 min. Thus, electrical signals produced by our engineered strain allow for rapid, continuous detection and quantification of thiosulfate.

To determine if our living electronic sensor can be rapidly diversified to respond to chemicals that affect vertebrate reproduction and health, we leveraged Fd switches that post-translationally gate ET in response to a chemical ligand (*21*, *26*). We replaced the native Fd in our I^+^C^+^O^+^ strain with a ligand-gated Fd that contains the estrogen receptor ligand-binding domain (*26*) and generated a Switch (S) strain (Fig. 3A), designated I^s^C^+^O^+^. We encapsulated the I^s^C^+^O^+^ and I^C42A^C^+^O^+^ strains into separate working electrodes, immersed them in the same anodic chamber in a 2-Encapsulated Working Electrodes (2-EWE) configuration (Fig. 3B), and added either DMSO or the endocrine disruptor 4-hydroxytamoxifen (4-HT). To quantify current changes induced by 4-HT, we calculated the percent difference in current of the I^s^C^+^O^+^ strain relative to the I^C42A^C^+^O^+^ strain 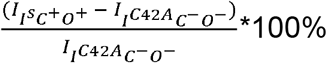. This comparison controls for any systemic environmental changes (*e.g.,* temperature, pH, carbon source) that affect the signal (*12*). Following 4-HT addition (12.5 μM), the I^s^C^+^O^+^ current increased within a few minutes (Fig. 3C). In contrast, the chemical used to dissolve 4-HT (DMSO) did not change the I^s^C^+^O^+^ current relative to I^C42A^C^+^O^+^. Comparison of the DMSO and 4-HT signals revealed that 4-HT was detected at 95% confidence in 7.8 min (Fig. 3D). Thus, the I^s^C^+^O^+^ strain responds to 4-HT as designed by producing enhanced current on the minute timescale. This living electronic sensor cuts the response time by a factor of ~4 compared with prior microbial bioelectronic sensors, which require between 0.5 to 5 h to respond to analytes (*8*–*12*, *27*–*29*).

**Fig. 3.**
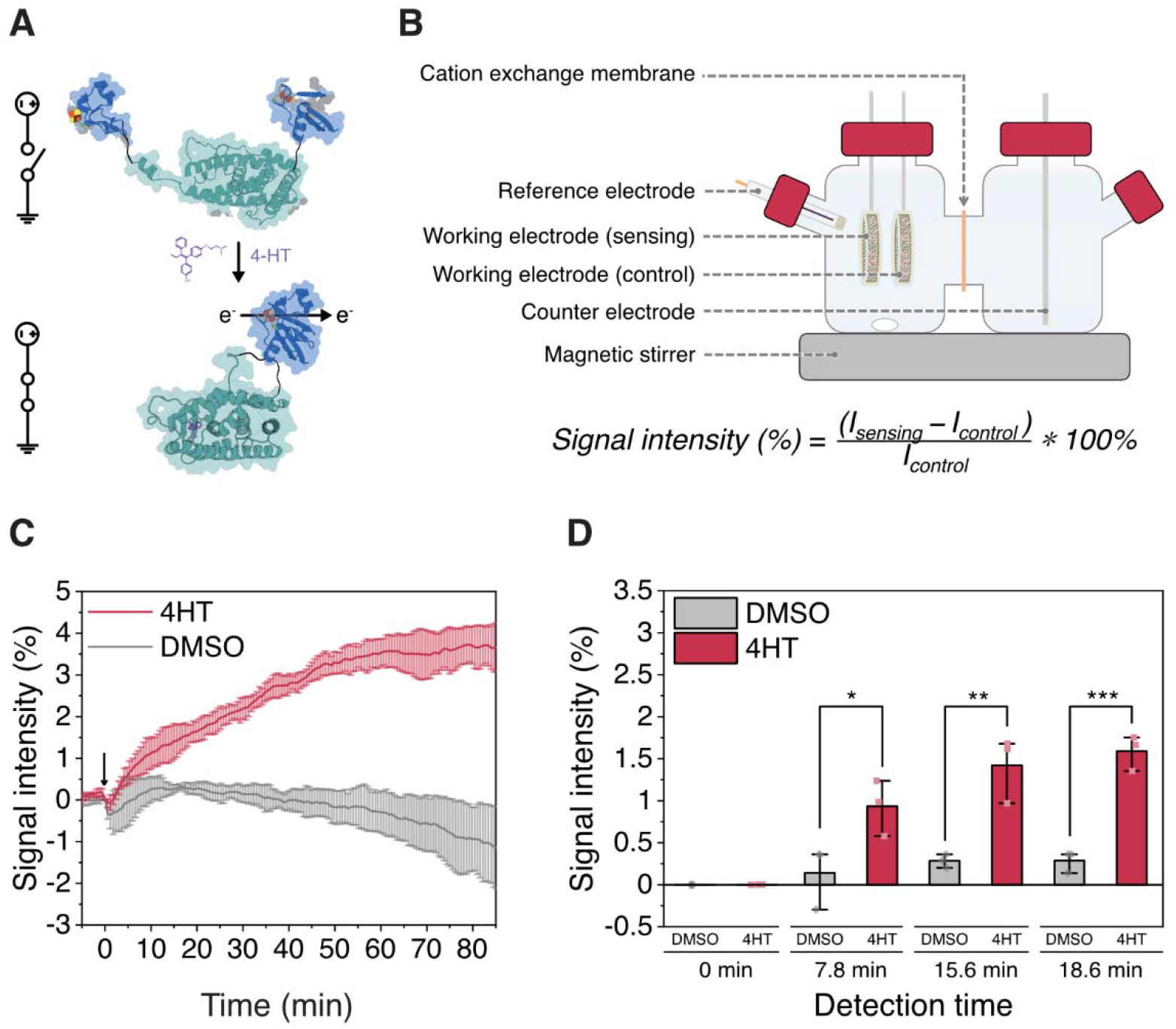
Living electronic sensors expressing an electrical protein switch yield rapid detection of an endocrine disruptor. **(A)** Schematic depicting a ligand-dependent Fd whose ET is regulated by 4-HT. **(B)** Schematic of the 2-EWE configured BES for 4-HT sensing, which contains two working electrodes: one encapsulating the I^S^C^+^O^+^strain, and the other containing the I^C42A^C^+^O^+^ strain. **(C)** Percent increase in current of I^S^C^+^O^+^ relative to I^C42A^C^+^O^+^ upon addition of DMSO or 4-HT in the 2-EWE configured BES. **(D)** Percent increase in current of I^S^C^+^O^+^ relative to I^C42A^C^+^O^+^ at different times following addition of DMSO or 4-HT. The times shown represent 4-HT detection with 95%(*), 99%(**), and 99.9%(***) confidence in 7.8 min (0.9% increase), 15.6 min (1.4% increase), and 18.6 min (1.6 % increase), respectively. Data represents the mean values with error bars representing one standard deviation (n = 3 biologically independent samples). P values were calculated using a one-way ANOVA with Tukey test.

To probe if our living electronic sensor functions in complex urban waterway samples, we tested our BES in riverine and marine samples spiked with thiosulfate or 4-HT. Water samples were collected from an urban beach (Galveston Beach) and two bayous (Buffalo Bayou and Brays Bayou) in the Houston Metro Area (Fig. 4A) that vary in pH, solution conductivity, and organic carbon content (Fig. 4B). We first tested thiosulfate sensing using I^+^C^+^O^+^ and I^C42A^C^+^O^+^ strains in a BES having a 2-EWE configuration. In all water samples, thiosulfate (10 mM) addition resulted in a significant increase (*p <* 0.05) in I^+^C^+^O^+^ current relative to the I^C42A^C^+^O^+^ current within 6.5 min (Fig. 4C), demonstrating our 2-EWE sensor functions robustly across urban water samples with different abiotic characteristics.

**Fig. 4.**
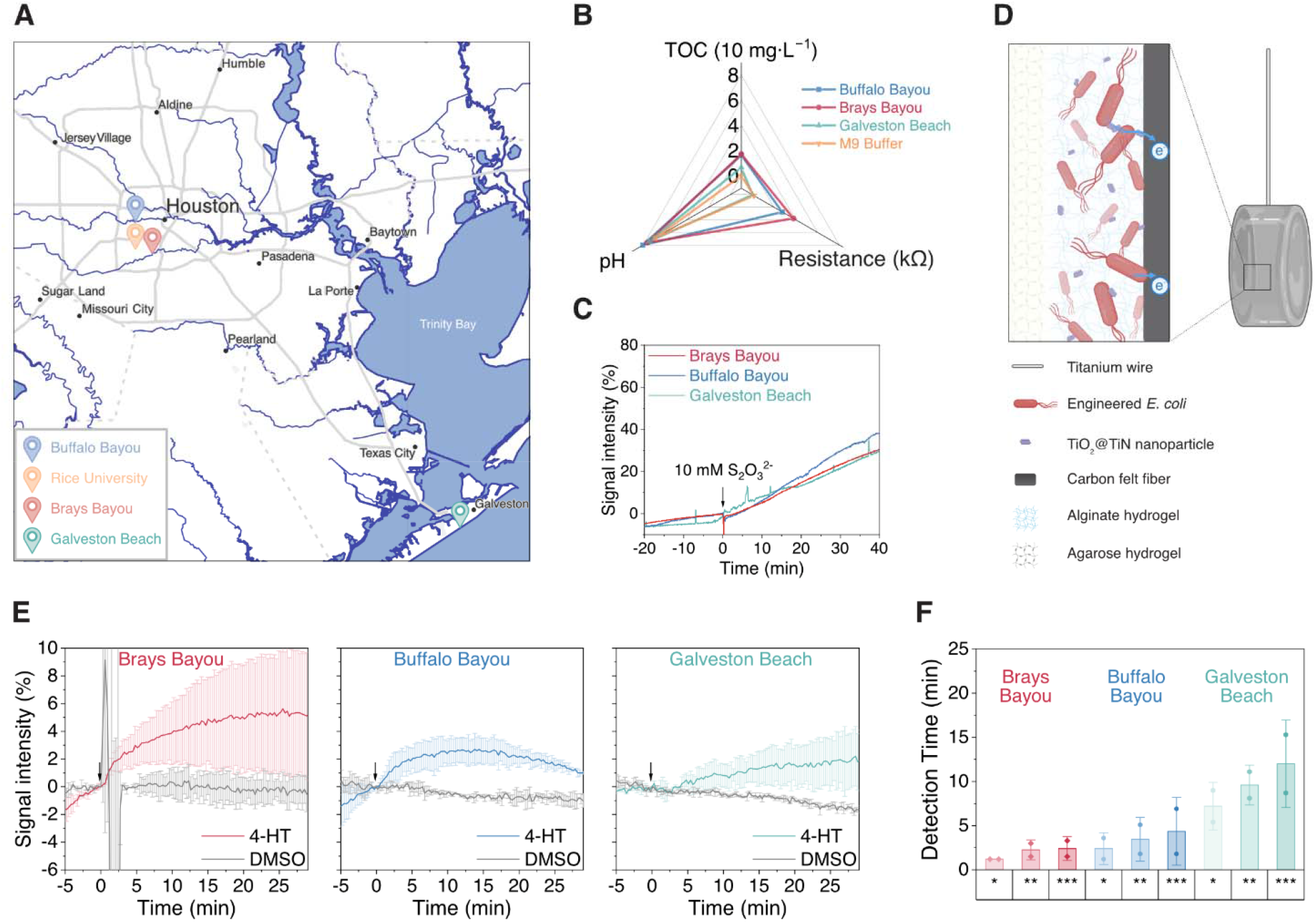
Living electronic sensors encapsulated with conductive nanoparticles enable rapid detection of pollutants in environmental samples. **(A)** Map of urban water sampling locations. **(B)** pH, solution resistance, and total organic carbon (TOC) measurements from each sample is compared with M9 medium. The pH ranged from 6.85 to 8.04, solution resistance ranged from 0.044 to 3.787 kΩ, and TOC ranged from 0 to 17.05 mg L^−1^. **(C)** The percent increase in current of I^+^C^+^O^+^ relative to I^C42A^C^+^O^+^ upon addition of thiosulfate in a 2-EWE configured BES using each environmental sample. Data represents the values from a single experiment in each environmental sample (n = 1 biologically independent samples). **(D)** Scheme illustrating the encapsulation with TiO_2_@TiN nanoparticles to enhance EET efficiency. **(E)** The percent increase in current of I^S^C^+^O^+^ relative to I^C42A^C^+^O^+^ upon addition of either 4HT or DMSO in a 2-EWE configured BES using each environmental sample. **(F)** Detection time for 4-HT with 95%(*), 99%(**), and 99.9%(***) confidence. Data in panel E and F represents the mean values with error bars representing standard deviation (n = 2 biologically independent samples). P values were calculated using a one-way ANOVA with Tukey test.

We next tested our living electronic sensors by adding 4-HT (12.5 μM) into the urban water samples using 2-EWEs containing the I^S^C^+^O^+^ and I^C42A^C^+^O^+^ strains. These water samples had poor conductivity (Fig. 4B) and abundant redox active compounds (Fig. S5), which pose additional challenges relative to laboratory media for bioelectronic sensing. To enhance current collection, we introduced a biocompatible and conductive TiO_2_@TiN nanocomposite into the encapsulation matrix (Fig. 4D), which increases the contact surface for strains to deliver electrons and facilitates electron transfer at the bacterial-electrode interface (*30*). Addition of nanoparticles led to an increase in the signal-to-noise with a higher steady-state current in the presence of 1 mM thiosulfate (Fig. S6), resulting in a faster response time.

When we used this novel encapsulation approach for urban water sensing, 4-HT caused immediate current increases with I^S^C^+^O^+^ across the sampling sites, while DMSO caused no deductible current increase (Fig. 4E). The response times for 4-HT detection were shortened to about 7 min with 95% confidence, with the fastest being less than 2 min. The response time required for 99.9% confidence was slightly longer (Fig. 4F), but always less than 15 min. Using a model of Fickian diffusion (*17*), we estimated the time required for 4-HT to penetrate the agarose layer and reach the bacteria (Fig S7). The agarose thickness varied between 1-3 mm, yielding a diffusional timescale between 1-8 min that agrees with our fastest response times. Thus, our living electronic sensor specifically detects analytes at environmentally-relevant concentrations and conditions with mass-transfer limited kinetics that are up to 10 times faster than the previous state-of-the-art (*8*–*12*, *27*–*29*, *31*).

This work describes three parallel innovations beyond previous work (*27*–*29*, *31*) required to achieve real-time sensing. First, our work introduces synthetic signal transduction using ET, in addition to phosphorylation (*32*) or proteolysis (*33*). This allows direct transmission of information and energy from biology to electronics (*33*). Second, the chemical gating of EET in this work is controlled post-translationally to enable rapid response times that are well suited for continuously monitoring transient chemical exposures in the environment. Third, we leveraged previous innovations in conductive nanomaterials (*30*) to improve the efficiency of EET within the encapsulation matrix, which amplified the signal and led to mass-transfer limited response times.

The modular approach used herein illustrates engineering principles and opportunities for how miniature bioelectronic devices can be created to enable real time multimodal sensing. In total, this pathway contains oxidoreductases from four different organisms across two domains of life that contain a total of twenty-nine cofactors. This demonstrates that electron transfer can be flexibly and extensively re-wired. This synthetic ET pathway could be adapted to respond to different analytes by inserting a wider range of ligand-binding domains and targeting different module components for switch design (*21*). To improve and customize this proof-of-concept for long-term environmental deployment, carbon sources and accessory chemicals can be incorporated within the synthetic encapsulation matrix, and these sensors can be incorporated into devices that self-power by scavenging energy present in the environment (*34*). Additionally, materials engineering can be used to optimize transmission of electrical signals at the abiotic-biotic interface. Small, deployable real-time bioelectronic sensors that can be distributed across different environmental locations will revolutionize our ability to monitor chemicals as they move through ecosystems informing smart sustainable practices in agriculture, mitigating the impacts of industrial waste release, and ensuring water security.

## ACKNOWLEDGMENTS

*E. coli* EW11 and the genes encoding FNR, and SIR were a gift from Dr. Pamela Silver (Harvard University). pSIM19 was a gift from Dr. Don Court (NIH-National Cancer Institute). pSS9 (Addgene plasmid #71655), pSS9-RNA (Addgene plasmid #71656), and pX2-Cas9 (Addgene plasmid #8581) were gifts from Dr. Ryan Gill (University of Colorado). Water sampling with help from Siliang Li (Rice University). TOC measurements with the help from Dr. Xiao Chen and Dr. Caroline Masiello (Rice University). We also thank Dr. Moshe Baruch (Rice University) for helping conceptualize the project.

## AUTHOR CONTRIBUTIONS

Contributions are noted in alphabetical order.

Conceptualization: CMAF, JTA, JJS, LS, XZ
Methodology: JTA, LS, XZ
Investigation: JTA, LS, XZ
Visualization: JTA, LS, XZ
Funding acquisition: CMAF, GNB, JJS
Project administration: CMAF, JJS
Supervision: CMAF, JJS
Writing – original draft: CMAF, JTA, JJS, LS, XZ
Writing – review & editing: CMAF, JTA, GNB, JJS, LS, XZ

## DATA AND MATERIALS AVAILABILITY

Genetic constructs will be made available in Addgene.

## SUPPLEMENTARY MATERIALS

Materials and Methods
Supplementary Text
Fig S1 – S7
Table S1 – S2
References (35 – 52)

