## Supplementary Materials for "Real-time environmental monitoring of contaminants using living electronic sensors"

**This PDF file includes:**

Materials and Methods
Supplementary Text
Figs. S1 to S7
Tables S1 to S2

**Materials and Methods**

Plasmid construction.

A list of all plasmids is in Table S1. To allow flexibility in use of different Fds, two plasmids were used to express the Input and Coupling modules. The first plasmid expresses FNR, SIR, SQR, and the Ccm system. The second plasmid expresses different Fds [Fd, Fd(C42A), or sFd-55-ER] as well as the *E. coli* MG1655 *fnr* and *pgl* gene cassettes. These latter genes were included to improve NADPH production and defects in anaerobic respiration. The parent strain EW11 and other BL21 derived strains respond to anoxic conditions in a manner distinct from K12 derived strains like MG1655 (*35*). One contributing factor is a nonsense mutation in the gene encoding the fumarate and nitrate reduction regulatory protein (FNR) (*36*). FNR is the global transcription factor responsible for activating expression of fumarate and nitrate reductases as well as many other genes involved in the transition from aerobic to anaerobic metabolism (*37*). BL21 derived strains also have a deletion of the *pgl* gene encoding 6-phosphogluconolactonase (35). This deletion limits flux through the oxidative branch of the pentose phosphate pathway requiring NADPH to be generated through TCA cycle (*38*), transhydrogenase (*39*) or potentially through one-carbon metabolism (*40*).

The Output module was chromosomally integrated. The genes encoding *Rhodobacter capsulatus* and *Geobacillus stearothermophilus* SQR were obtained as Gblocks (Integrated DNA Technologies) and cloned using Golden Gate DNA assembly (41) into pSAC01 (*21*) to create pSAC01_SQR1 and pSAC01_SQR2, respectively. A constitutively expressed *ccm* operon from *E. coli* was amplified from pM0640 (*20*) and cloned into pSAC01, pSAC01_SQR1, and pSAC01_SQR2 using Gibson DNA assembly (42) with unique nucleotide sequences (UNS3 and UNS4) (43) to generate p(e-)nzymes_NC, p(e-)nzymes, and pJA036, respectively. The *fnr* and *pgl* gene cassettes were PCR amplified from genomic *E. coli* MG1655 DNA and cloned into pFd007/lacI (*21*), pFd007_C42A/lacI (*21*), and pBW014 (*26*) using Golden Gate to generate pFd007/lacI/fnr/pgl, pJA035, and pJA040, respectively. To facilitate chromosomal integration of the Output module, a T7-lac driven version of the *cymA-mtrCAB* operon from *Shewanella oneidensis* MR-1 was PCR amplified from pI5049 (*19*) and cloned into pSS9 (*44*) using restriction enzyme cloning with KpnI and SpeI to generate pSS9:cymAmtrCAB. pSIM19 was from Dr. Don Court. pSS9 (Addgene plasmid #71655), pSS9-RNA (Addgene plasmid #71656), and pX2-Cas9 (Addgene plasmid #8581) were from Dr. Ryan Gill. All plasmids were sequence verified using Sanger DNA sequencing.

Strains.

A list of all strains is in Table S2. *Escherichia coli* XL1-Blue (Stratgene) was used for all plasmid construction and amplification. All other experiments were performed using the *E. coli* EW11 [B F- dcm ompT hsdS($r_B^-$ $m_B^-$) gal λ(DE3) Δ*cysI fpr ydbK hcr yeaX hcaD frdB hycE hyaB hybC hyfG*] (*22*) or *E. coli* EW11-JA01 [B F- dcm ompT hsdS($r_B^-$ $m_B^-$) gal λ(DE3) Δ*cysI fpr ydbK hcr yeaX hcaD frdB hycE hyaB hybC hyfG* ss9::T7-*cymA-mtrCAB*], which was from Dr. Pam Silver.

To build *E. coli* EW11-JA01, CRISPR-recombineering (44) was used to integrate the *cymA-mtrCAB* operon under control of the T7-lac promoter at safe-site 9 (SS9) (*44*) in the *E. coli* EW11 genome. EW11 was made CRISPR-recombineering ready by transformation with pSIM19 (*45*) and pX2-Cas9 (44). After selection on lysogeny broth (LB) agar plates with 50 µg/mL kanamycin and 100 ug/mL streptomycin at 30 °C, a fresh colony was picked and grown in 3 mL of LB with 50 µg/mL kanamycin and 100 µg/mL streptomycin at 30 °C for 18 h. Cultures were diluted 1:100 into LB medium (50 mL) containing 50 µg/mL kanamycin and 100 µg/mL streptomycin and grown at 30 °C to exponential phase. The λ red recombination machinery was induced by incubating at 42°C for 15 min. Cells were then concentrated (500x) and made electrocompetent by centrifugation (6000 x g) and washed with 10% glycerol four times.

The T7-lac driven *cymA-mtrCAB* operon flanked by 100 bp was amplified from pSS9:cymAmtrCAB using primers JA307 (CCTGAGCTTGATCCTACAC) and JA308 (5'-GACAGGATGATTACATAAATAATAGTG-3'). This linear, double-stranded DNA fragment was isolated by agarose gel extraction to obtain a substrate for CRISPR-recombineering. CRISPR-recombineering ready cells (50 µL) were electroporated with 100 ng of pSS9-gRNA and 500 ng of the T7-lac driven *cymA-mtrCAB* operon flanked by 100 bp of SS9 homology and recovered for 3 h at 30 °C in 1 mL of LB containing 0.2% arabinose, which induced Cas9 expression. Cells were then plated onto LB agar plates supplemented with 0.2% arabinose, 25 µg/mL kanamycin, and 50 µg/mL ampicillin and grown for 18 h at 37 °C.

Integration was verified using PCR amplification of genomic DNA from the EW11 parent strain and EW11-JA01 strain purified using a Wizard® Genomic DNA Purification Kit (Promega). The SS9 locus was amplified using primers JA299 (5'-CATGTCGTCAAAATGTTG-3') and JA300 (5'-TTTGATGTTAACGTTGCAGA-3'). This ~6.7 kb band was agarose gel purified and sequence verified using Sanger DNA sequencing with primers JA299 and JA300.

Media and cell growth conditions.

All molecular biology was performed in LB medium. For growth and sulfide production assays evaluating Input and Coupling module functions, cells were grown in M9c or M9sa (*21*). Colonies from freshly streaked LB agar plates were used to inoculate 1 mL of M9c with 100 µg/mL streptomycin and 34 µg/mL chloramphenicol in 2 mL 96-well polypropylene plates (USA Scientific No. 1896-2110) and were grown at 37 °C shaking at 250 rpm for 18 h. An aliquot of this culture (1 µL) was used to inoculate M9sa (100 µL) containing 100 µg/mL streptomycin, 34 µg/mL chloramphenicol, 10 µM IPTG, and varying aTc as noted in a 96-well polystyrene plate (Corning No. 3595). Cells were then grown at 37 °C shaking at 250 rpm for 24 h.

For evaluation of Output module expression, EET, and electrochemical assays, biomass was generated by inoculating 2xYT media supplemented with 200 µM 2-aminolevulinic acid, 1x trace minerals, 100 µg/mL streptomycin, 34 µg/mL chloramphenicol with 1:100 v/v of a culture grown overnight in LB with antibiotics. Cultures were grown aerobically at 37 °C with 250 rpm shaking until they reached exponential phase ($OD_{600}$ = 0.5-0.6). To induce the expression of the *cymA-mtrCAB* operon expression, varying IPTG and aTc were added as noted and cultures were grown at 30 °C with 250 rpm shaking for 18 h. The 100x trace mineral stock solution (pH 7.0) contained: 7.85 mM $C_6H_9NO_3Na_3$, 12.17 mM $MgSO_4·7H_2O$, 2.96 mM $MnSO_4·H_2O$, 17.11 mM NaCl, 0.36 mM $FeSO_4·7H_2O$, 0.68 mM $CaCl_2·2H_2O$, 0.42 mM $CoCl_2·6H_2O$, 0.95 mM $ZnCl_2$, 0.040 mM $CuSO_4·5H_2O$, 0.021 mM $AlK(SO_4)_2·12H_2O$, 0.016 mM $H_3BO_3$, 0.010 mM $Na_2MoO_4·2H_2O$, 0.010 mM $NiCl_2·6H_2O$, and 0.076 mM $Na_2WO_4·2H_2O$.

Sulfide oxidation measurement.

Sulfide oxidation by cells expressing SQRs was monitored using a sulfide selective microsensor (SULF-MR, Unisense A/S) connected to a four-channel multiprobe micrometer (Unisense A/S). To calibrate the microsensor an 8-point standard curve was generated using a freshly prepared solution of sodium sulfide in 1x M9 salts prior to measurements. Cells grown to stationary phase in M9c were washed and resuspended to an $OD_{600}$ = 0.5 in 1x M9 salts. This suspension (4 mL) was transferred to a respiration chamber (Unisense A/S). Following a 2 min baseline measurement, 500 μM sodium sulfide was injected into the chamber and sulfide was monitored every 1 s for 15 min. The mean and standard deviation of the sulfide concentration for biologically independent samples (n = 3) are reported. First order rates were calculated using linear regression.

Bioelectrochemical analysis of Output module EET.

To characterize EET by the $I^{C42A}C^-O^+$ and $I^{C42A}C^-O^-$ strains, bioelectrochemistry measurements were carried out in dual-chamber (150 mL in volume per chamber) bioelectrochemical reactors (Adams & Chittenden Scientific Glass) using a VSP-300 potentiostat (BioLogic). The anodic chamber contained an Ag/AgCl reference electrode (3 M KCl, CHI111, CH Instruments) and a 6.35-mm-thick graphite felt working electrode with a 16-mm radius (Alfa Aesar). The cathodic chamber contained a 0.5-mm radius titanium wire as the counter electrode (Alfa Aesar). The two chambers were separated by a cation exchange membrane (CMI7000, Membranes International). Each chamber contained ~125 mL M9 buffer and 0.2% glucose, unless otherwise indicated. Both anodic and cathodic chambers were kept at 30 °C by placing the reactors in an incubator.

To characterize current production by the $I^{C42A}C^-O^+$ and $I^{C42A}C^-O^-$ strains, controlled potential chronoamperometry was carried out under anoxic conditions. To maintain anoxic conditions, reactors were continuously purged with pure $N_2$ gas by inserting a needle into the M9 buffer for the duration of experiment. The working electrode was held at +0.42 $V_{SHE}$. Once the current stabilized, the washed strains were inoculated into the anodic chamber with a final $OD_{600}$ of 0.5. The medium in the electrochemical chamber was mixed with a magnetic stir bar at 250 rpm mixing rate for the course of the experiment. The average current over every 36 s was recorded, and results are representative of three independent experiments, unless otherwise indicated.

Bioelectrochemical analysis of thiosulfate and 4-HT effects on EET.

For bioelectronic sensing of both thiosulfate and 4-HT, bioelectrochemistry measurements were carried out in water-jacketed dual-chamber (125 mL in volume per chamber) bioelectrochemical reactors (Adams & Chittenden Scientific Glass) using a VSP-300 potentiostat (BioLogic). These measurements used the same Ag/AgCl reference electrodes (3 M KCl, CHI111, CH Instruments) and 0.5-mm radius titanium wire counter electrodes (Alfa Aesar), but smaller working electrodes (6.35-mm-thick graphite felt with a 10.5-mm radius) were used to fit the smaller reactor chambers. Each chamber contained ~115 mL M9 buffer and 0.2 % glucose, unless otherwise indicated. Both anodic and cathodic chambers were kept at 30 °C by connecting the water-jackets to an ECO E4S heating circulator (Lauda-Brinkmann).

To characterize current production by $I^+C^+O^+$ and $I^{C42A}C^+O^+$ strains for thiosulfate sensing under laboratory conditions, chronoamperometry was carried out as described above. After strains were injected and currents stabilized, an increasing concentration of sodium thiosulfate was injected into the reactor over the course of 40 min. The average current over every 36 s was recorded, and results are representative of three independent experiments, unless otherwise indicated. Current after 5 min and 30 min from each injection were used for linear analysis (OriginPro 2021, OriginLab Corporation) between the thiosulfate concentration and current response.

For all 4-HT sensing, and thiosulfate sensing within environmental samples, we introduced a novel 2-Encapsulated Working Electrodes (2-EWE) system instead of the previous single working electrode system. The 2-EWE system included two types of strains ($I^SC^+O^+$ and $I^{C42A}C^+O^+$ strains for 4-HT sensing, or $I^+C^+O^+$ and $I^{C42A}C^+O^+$ strains for thiosulfate sensing) encapsulated separately with the working electrodes in the same reactor chamber, which generated two current signals under the same environmental conditions simultaneously by connecting to two potentiostat channels. The potentiostat was operated under a ‘counter electrode to ground’ mode by connecting one shared counter electrode which was grounded, one shared reference electrode, and two individual working electrodes. During the test, there were also some changes to mimic the practical biosensing scenario: river or marine water was filtered with 0.2 μm filter, and then used in the reactors as electrolyte without any additives; the needle for purging $N_2$ gas was lifted just above the liquid level after working electrodes were introduced to minimize the disturbance from purging; magnetic stirring was also stopped for the same purpose; for 4-HT or DMSO sensing, 10 mM sodium thiosulfate and 0.2% glucose were also include in the M9 buffer solution. For increased resolution, the average current over every 3.6 s was recorded, and results are representative of three independent experiments, unless otherwise indicated.

As previously described, the 2-EWE system allows measurement of current signals from two different strains at the same time and condition. By comparing the difference between these two signals, any other systemic effects (such as changes in temperature, pH, carbon source or else) can be excluded, such that the signal represents our designed ET, *i.e.*, the response to only 4-HT or thiosulfate. Currents are reported as the percent change between the sensing ($I^+C^+O^+$ or $I^SC^+O^+$) and the control ($I^{C42A}C^+O^+$) strains:

$$\text{Signal intensity (\%)} = (I_{sensing} - I_{control}) / I_{control} * 100\%$$

<u>Cell encapsulation.</u>

Concentrated washed strains ($OD_{600}$ = 40) and sodium alginate solution (3.9 wt%, in M9) were mixed at a 1:1 ratio on ice. The mixture (1 mL) was applied to a carbon felt electrode (10.5 mm radius) at room temperature, and it was solidified by immersing into $CaCl_2$ aqueous solution (3 wt%) for 30 s. The residual chemicals on the hydrogel-electrode surface were washed with M9 medium. Subsequently, the formed hydrogel was covered with 1 wt% agarose to provide mechanical support.

The $TiO_2$@TiN nanocomposite was synthesized as previously reported (*30*). In brief, $TiO_2$ nanoparticles (P25, Alfa Aesar) were heated at 900 °C in a tube furnace (KTL1400, Nanjing University Instrument Plant, China) in an ammonia atmosphere for 2 h. When mixing with the strains and sodium alginate solution, 1 mg $mL^{-1}$ $TiO_2$@TiN nanocomposite was added.

Water sampling and characterization.

The environmental samples were collected from the Houston area and stored at 4 °C. Before the measurement, all environmental samples were filtered with 0.2 μm sterile membranes to remove solids and microorganisms. The pH of each sample was measured by a pH meter (F20, Mettler Toledo). Resistance was measured by electrochemical impedance spectroscopy (EIS) using a three-electrode system in 125 mL dual-chamber reactors, with an amplitude of 5 mV over a frequency range of 100 kHz to 0.01 Hz at open circuit potential. The results were analyzed with ZView 3.5b (Scribner Associates Inc). Total organic carbon (TOC) analysis was conducted on a TOC-vcsh analyzer (Shimadzu). Cyclic voltammetry was performed using a VSP-300 potentiostat (BioLogic) and scanned from -0.8 V to 0.6 V vs Ag/AgCl, scan rate is 10 mV/s.

Cytochrome expression and function analysis.

After growth in 96-well plates, 150 μL of culture was pelleted in white, U-bottom 96-well polystyrene plates (Corning No. 3917). Prior to pelleting, the cells were washed and resuspended in anoxic M9 minimal medium supplemented with 100 mM sodium lactate. Additionally, 3 mg/mL $WO_3$ nanoparticles were added and incubated under anoxic conditions for 6 h at 30 °C prior to pelleting to evaluate EET. To evaluate redness and blueness for each assay, the plates containing the cell pellets were scanned using a desktop scanner. Regions of interest from each pellet were identified using the Matlab (R2018a) ImageProcessing Toolbox (MathWorks). The relative red intensity was calculated by taking the ratio intensity in the red channel ($I_{red}$) to the grayscale intensity ($I_{gray}$), red intensity = ($I_{red}$)/($I_{gray}$). The relative blue intensity was calculated by taking the ratio of the intensity in the red channel ($I_{blue}$) to the grayscale intensity ($I_{gray}$), blue intensity = ($I_{blue}$)/($I_{gray}$). For blue intensity analysis, $WO_3$ oxide nanoparticles were synthesized as described (46). In brief, 0.85 g $Na_2WO_4 \cdot 2H_2O$ and 0.29 g NaCl were dissolved in 20 mL dd$H_2O$, then adjusted the pH to 2.0 with 3M HCl. The solution was transferred into a hydrothermal reactor and heated at 180 °C for 7 h. $WO_3$ nanoparticles were then harvested by washing with dd$H_2O$ until the supernatant reached pH 7.0, then collected the solid with filtering through a 0.45 μm membrane.

Sulfane sulfur analysis.

To measure intracellular sulfur accumulation following sulfide oxidation by SQR, a fluorescent probe for sulfane sulfur was used as described (*25*). Cells were washed 2x in 50 mM HEPES (pH 7.4) and resuspended to $OD_{600}$ = 2 in HEPES buffer containing 10 μM SSP4 (3',6'-Di(O-thiosalicyl)fluorescein, Dojindo Laboratories) and 0.5 mM dodecyltrimethylammonium bromide. Cells were incubated for 15 min at 37 °C in the dark with gentle shaking (150 rpm) and washed 2x with HEPES buffer. $OD_{600}$ and fluorescence intensity ($\lambda_{excitation}$= 482 nm; bandwidth = 20 nm, $\lambda_{emission}$= 515 nm; bandwidth = 20 nm) were quantified using a Spark microplate reader (Tecan). The mean and standard deviation of the OD normalized fluorescence intensity for biologically independent samples (n = 3) are reported.

Hydrogen sulfide production.

Evolution of $H_2S(g)$ from cultures was monitored using a semi-quantitative lead acetate filter paper assay that monitors the formation of insoluble PbS pigment in 96-well plates (47). Whatman No.1 filter paper was cut to fit inside the lid of a 96-well plate (Costar No. 3526). A 2% lead acetate solution in water was prepared and particulates were removed by filtration through a 0.22 µm filter. The cut filter paper was then soaked in 2% lead acetate for 30 min. Filter paper was removed and allowed to air dry overnight. Dried filter papers were applied to the inside of the 96-well plate prior to growth experiments. Following incubation with cultures in the plates, the filter paper was scanned and the intensity of the gray channel was quantified using the FIJI image processing package with the ReadPlate3 plugin (https://sites.imagej.net/ReadPlate/plugins/). The mean and standard deviation of the gray channel intensity for biologically independent samples (n = 3) are reported.

Calculation of diffusional timescale

To estimate the time required for analytes, such as 4-HT or thiosulfate, to reach the encapsulated *E. coli*, we calculated the diffusional timescale following the model of Liu *et al. (17)*. This model assumes an initial concentration of analyte, $I_o$, is separated by a cell-free hydrogel layer of thickness *L* from the bacteria (Fig S6). This model determines the time for the analyte concentration to reach *K*, the minimum analyte concentration needed to generate a response, at position *L* using an approximate solution to Fick's Law in 1D:

$$t_{diffuse} \approx \frac{4}{9} \frac{1}{\left(\frac{I_o}{K} - 1\right)^{0.56}} \frac{L^2}{D_g}$$

Since the diffusion constant of small molecules in agarose is typically ~95% that of in water *(48)* and the diffusion constant of thiosulfate in water ($D_{water}$) is 6.6 x $10^{-10}$ $m^2$ $s^{-1}$ *(49)*, we set $D_g = D_{water}$ *0.95 = 6.3 x $10^{-10}$ $m^2$ $s^{-1}$. We set K and $I_o$ equal to 0.1 mM (Fig 2) and 10 mM, respectively. Due to our fabrication process, *L* was not stringently controlled and varied between 1-3 mm. Since $t_{diffuse}$ depends on $L^2$, this variability translates into a significant variation in the critical diffusion timescale, yielding an estimated $t_{diffuse}$ between 1-8 min.

Statistics.

All reported P values were obtained using two-tailed, independent t-tests or one-way ANOVA using Tukey's test as noted. Sample sizes were in accordance with community standards.

### Supplementary Text

Evaluation of sulfur source on assimilation and $H_2S$ evolution.

*E. coli* assimilates inorganic sulfur sources (*e.g.,* sulfate, sulfite, thiosulfate) that vary in energetic costs for fixation into L-cysteine (Fig. s3A). Sulfur metabolism is tightly regulated in *E. coli* to limit the overproduction of L-cysteine, maintaining a total sulfur-atom concentration inside *E. coli* of ~0.13 M (50). This regulation is vital as at high concentrations L-cysteine can cause DNA damage through the Fenton reaction (51) and its degradation to sulfide can lead to cytochrome *bo* oxidase inhibition (50). Sulfate assimilation requires 3 mol of ATP and 4 mol of NADPH/mol of $SO_4^{2-}$ assimilated to yield one mol of L-cysteine. In contrast, thiosulfate assimilation is energetically less expensive yielding 2 mol of L-cysteine/mol of $S_2O_3^{2-}$, while only requiring only 1 mol of ATP and 4 mol of NADPH (Fig s3A). Thiosulfate, sulfate, or sulfite enter the cytosol through the ATP-dependent anion permease consisting of CysAUW and the periplasmic binding protein CysP or Sbp or the proton-dependent symporter CysZ. Sulfate proceeds through the canonical sulfate assimilation pathway becoming activated by ATP sulfurylase (CysDN) and APS kinase (CysC) using 2 ATP and reduced by PAPS reductase (CysH) using 1 NADPH to generate sulfite. Sulfite serves as the substrate for SIR using 3 NADPH to generate sulfide with FNR and Fd delivering these electrons to SIR in our engineered strains. Sulfide is then attached to the carbon skeleton *O*-acetyl-sertine (OAS) to generate L-cysteine by cysteine synthase (CysK). In contrast, thiosulfate is initially attached to the OAS skeleton by cysteine synthase B (CysM) to form *S*-sulfocysteine which undergoes reductive cleavage by glutaredoxin using 1 NADPH to form L-cysteine and sulfite (52). Sulfite can be assimilated into an additional L-cysteine through SIR and CysK. OAS is generated from L-serine by serine acetyltransferase (CysE) and flux is regulated post-translationally through feedback inhibition by L-cysteine. When OAS becomes limiting, sulfide accumulates in the cell and negatively regulates expression of the sulfate assimilation pathway by inhibiting the CysB regulon. If sulfide or thiosulfate becomes limiting, *N*-acetylserine (NAS) forms from OAS and positively regulates expression of the sulfate assimilation pathway by activating the CysB regulon.

To determine how Fd expression influences sulfide production from these different sulfur sources, we monitored growth (Fig. s3B) and $H_2S$ evolution (Fig. s3C) from $I^+C^-O^-$ cultures grown in medium containing these anions. In sulfate and sulfite media, cells displayed Fd-dependent growth that increased with aTc concentration. In contrast, $H_2S$ evolved from these cultures only at intermediate aTc concentrations. Cells grown with thiosulfate grew at all aTc concentrations. Additionally, thiosulfate cultures evolved similar levels of $H_2S$ at all aTc concentrations. The differences in $H_2S$ evolution at high aTc concentrations was interpreted to be caused by these differences in regulatory topologies with sulfate assimilation structured as an incoherent feedforward loop which can lead to band-pass filter behavior (53, 54) while thiosulfate assimilation is regulated by a post-translational negative feedback loop.

**Fig. S1. Effect of Output module expression on cytochrome levels. (A)** Red color intensity values of $I^{C42A}C^{-}O^{+}$ (circle) or $I^{C42A}C^{-}O^{-}$ (square) cell pellets following aerobic growth in 2xYT medium containing varying amounts of IPTG, which induces expression of CymA-MtrCAB. **(B)** Blue color intensity values of $I^{C42A}C^{-}O^{+}$ (circle) or $I^{C42A}C^{-}O^{-}$ (square) in minimal media containing lactate and electrochromic $WO_3$ nanoparticles that change from white to blue when reduced by microbes that present EET. **(C)** Cell density ($OD_{600}$) of $I^{C42A}C^{-}O^{+}$ (circle) or $I^{C42A}C^{-}O^{-}$ (square) grown in M9 minimal medium containing varying amounts of IPTG. Growth of $I^{C42A}C^{-}O^{+}$ was significantly decreased at >10 μM IPTG (12.5 μM: $p = 6.55 \times 10^{-3}$, 25 μM: $p = 7.30 \times 10^{-6}$, 50 μM: $p = 3.87 \times 10^{-4}$, 100 μM: $p = 4.65 \times 10^{-6}$, 200 μM: $p = 5.89 \times 10^{-5}$). Data represents the mean values with error bars representing one standard deviation (n = 3 biologically independent samples). P values were obtained using two-tailed, independent t-tests.

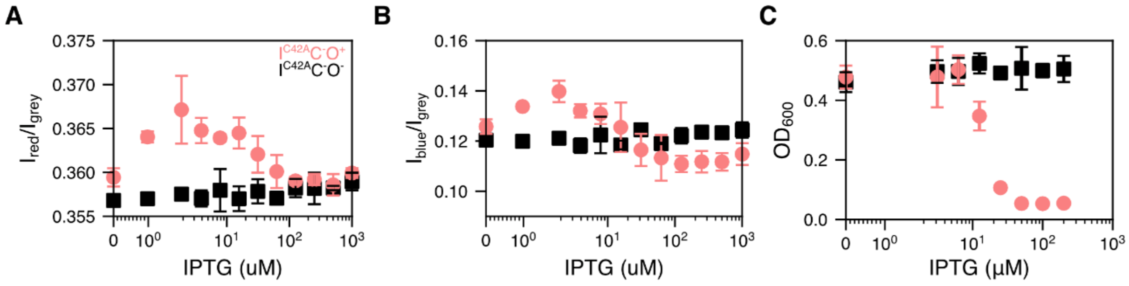

**Fig. S2. Sulfane sulfur accumulation in SQR expressing cells.** Relative fluorescence of cells treated with the sulfane sulfur probe SSP4. The fluorescence from cells expressing *Gs*-SQR and *Rc*-SQR was significantly higher than cells transformed with an empty vector (EV) ($p$ = 4.89 x $10^{-3}$ and $p$ = 3.14 x $10^{-3}$, respectively). Fluorescence from cells expressing *Gs*-SQR and *Rc*-SQR were not significantly different. Error bars represent one standard deviation (n = 3 biologically independent samples) with individual samples shown as white circles and bars heights representing the average. P values were obtained using two-tailed, independent t-tests.

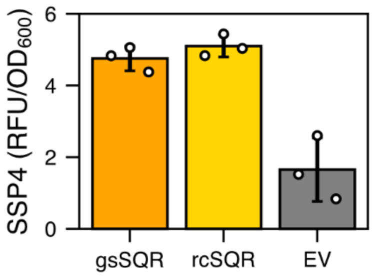

**Fig. S3. Impact of sulfur source on sulfide evolution from *E. coli* EW11. (A)** A schematic of sulfur metabolism and regulation in *E. coli* (*yellow*) and the redox coupling of the Input module (*blue*) with this pathway. **(B)** PbS accumulation and **(C)** optical density of the $I^{+}C^{-}O^{-}$ strain containing a vector for expressing Fd after 24 h in M9sa medium containing 2 mM of sulfate, sulfite, or thiosulfate and varying amounts of aTc to control Fd expression. Optical density in sulfate and sulfite containing media were significantly lower than thiosulfate containing media when <7.8125 nM aTc was added ($p < 0.01$). **(D)** PbS accumulation and **(E)** optical density after 24 h in M9sa medium containing varying amounts of thiosulfate and varying amounts of aTc to control Fd expression. In media with ≥0.25 mM thiosulfate, optical densities were not significantly different regardless of aTc concentration ($p > 0.01$). For panels B-E, symbols and error bars represent the mean and standard deviation, respectively (n = 3 biologically independent samples). P values were obtained using two-tailed, independent t-tests.

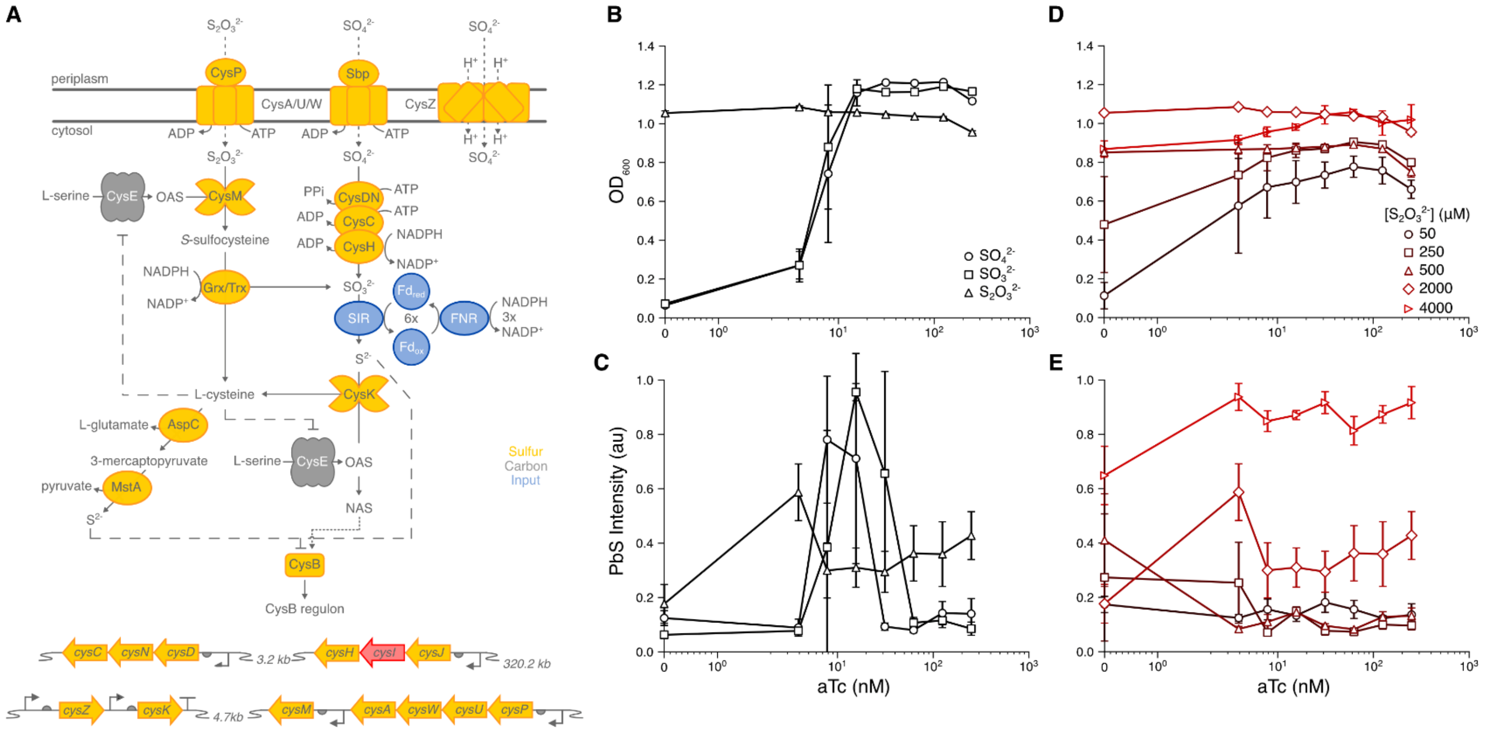

**Fig. S4. Planktonic cells present a small, noisy current response to thiosulfate.** The chronoamperometric response of planktonic $I^{+}C^{+}O^{+}$ and $I^{C42A}C^{+}O^{+}$ cells in a bioelectrochemical reactor. Arrows indicate the addition of thiosulfate to varying concentrations. Data represents the mean values with error bars representing one standard deviation (n = 3 biologically independent samples).

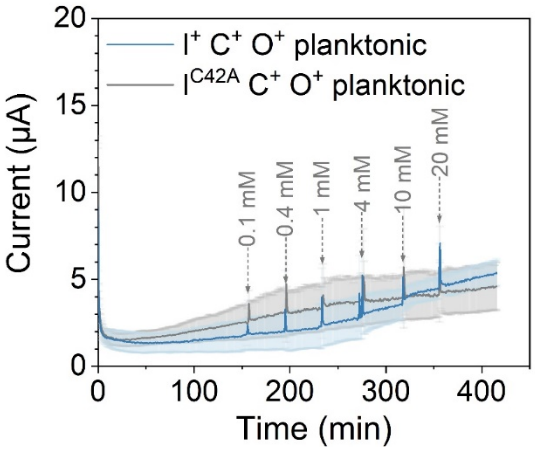

**Fig. S5. Cyclic voltammetry analysis of environmental samples. (A)** Each environmental sample shows multiple pairs of redox peaks, indicating abundant redox active chemicals exist which might interfere with 4-HT sensing. **(B)** Environmental samples supplemented with 0.2% glucose show no changes to their voltammograms. All CVs were measured at a scan rate of 10 mV/s.

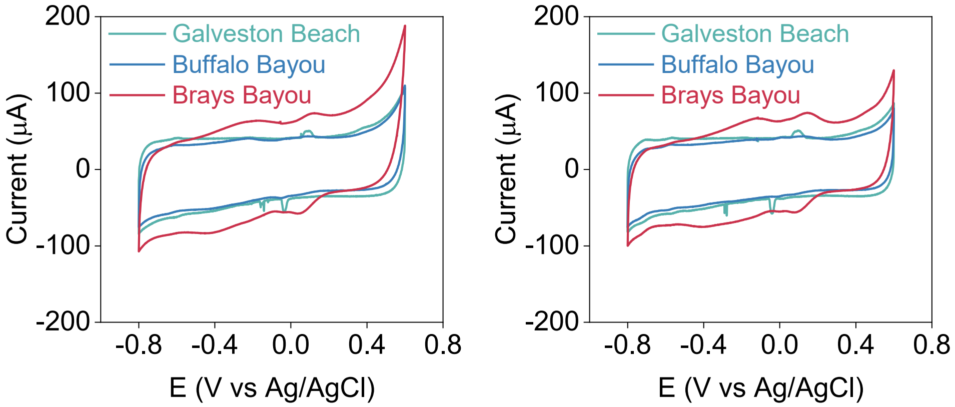

**Fig. S6. Addition of $TiO_2$@TiN nanoparticles enables more current collection. (A)** Chronoamperometry and **(B)** current of $I^+C^+O^+$ strain encapsulated in an alginate-agarose hydrogel with and without $TiO_2$@TiN nanoparticles upon addition of 1 mM thiosulfate (arrow). The strains encapsulated with nanoparticles respond to thiosulfate more rapidly and with a higher steady-state level. Data represents the mean values with error bars representing one standard deviation (n = 3 biologically independent samples). P values were calculated using a one-way ANOVA with Tukey test.

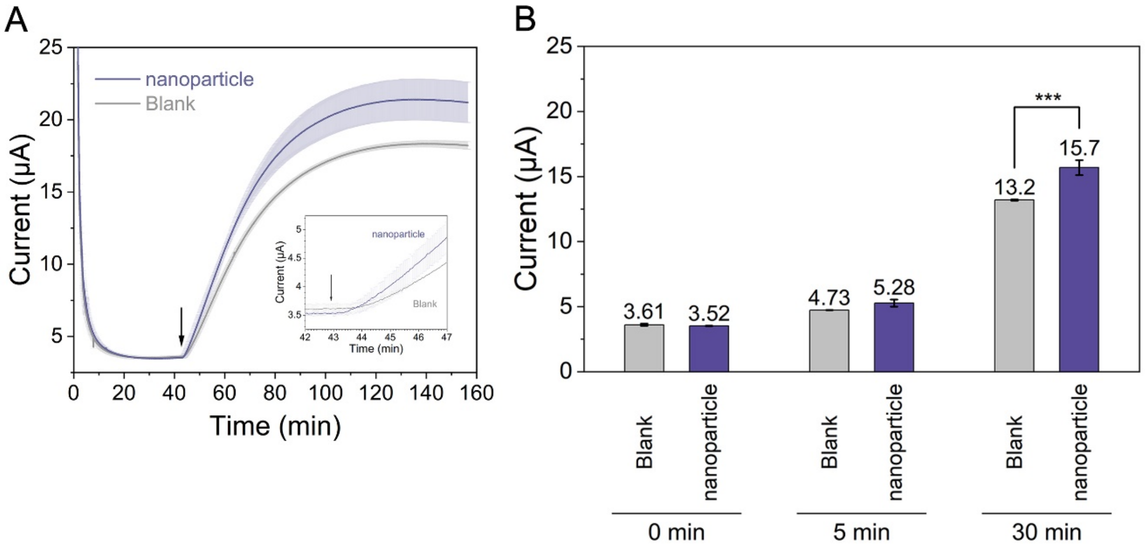

**Fig. S7. Simplified 1D geometry for calculation of diffusion timescales for response of living bioelectronic sensor.** Schematic of analyte diffusion from bulk solution through the agarose layer to cells embedded in the hydrogel on electrode surface.

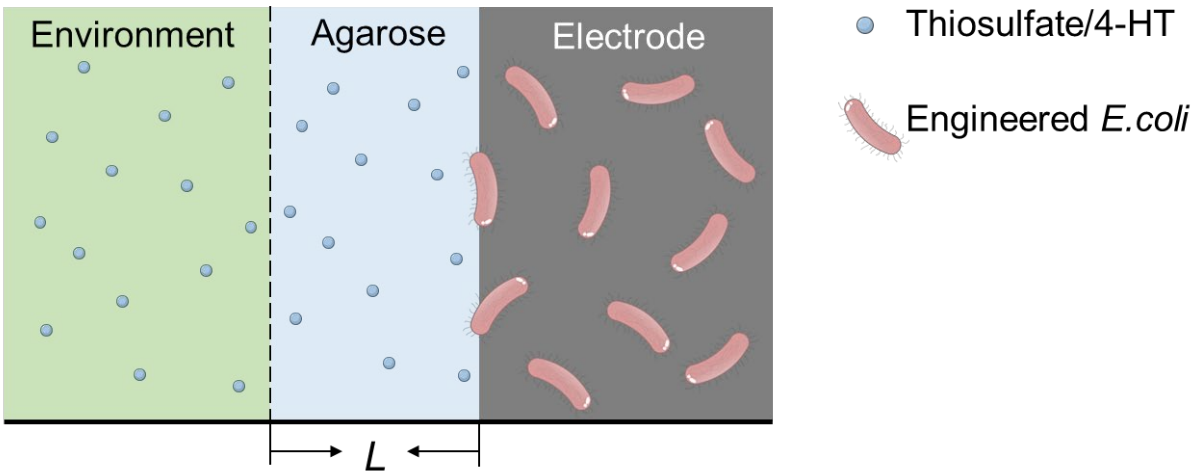

**Table S1. List of plasmids.** For each plasmid, the protein components expressed are noted.

| Plasmid Name | Description | Addgene ID |
|---|---|---|
| pSAC01 | FNR/SIR expressing plasmid | 131826 |
| pSAC01_SQR1 | FNR/SIR/rcSQR expressing plasmid | *To be submitted* |
| pSAC01_SQR2 | FNR/SIR/gsSQR expressing plasmid | *To be submitted* |
| p(e-)nzymes_NC | Ccm/FNR/SIR expressing plasmid | *To be submitted* |
| p(e-)nzymes | Ccm/FNR/SIR/gsSQR expressing plasmid | *To be submitted* |
| p(e-)nzymes_2 | Ccm/FNR/SIR/rcSQR expressing plasmid | *To be submitted* |
| pFd007/lacI | Fd expressing plasmid | 131832 |
| pFd007_C42A/lacI | Fd(C42A) expressing plasmid | 131831 |
| pBW014 | sFd-55-ER expressing plasmid | 131817 |
| pFd007/lacI/fnr/pgl | Fd/Fnr/Pgl expressing plasmid | *To be submitted* |
| pFd007_C42A/lacI/fnr/pgl | Fd(C42A)/Fnr/Pgl expressing plasmid | *To be submitted* |
| pERA007.55/lacI/fnr/pgl | sFd-55-ER/Fnr/Pgl expressing plasmid | *To be submitted* |
| pSIM19 | lambda red recombinase expressing plasmid | *To be submitted* |
| pX2-Cas9 | Cas9 expressing plasmid | 85811 |
| pSS9-RNA | gRNA expressing plasmid | 71656 |
| pSS9 | ss9 homology arm containing plasmid | 71655 |
| pSS9:cymAmtrCAB | ss9 homology arm flanking cymA-mtrCAB operon | *To be submitted* |

**Table S2. List of strains.** In total, three different strains were used, including XL1-Blue, EW11, and EW11-JA01. To create strains that express different combinations of the **I**, **C**, and **O** module components, strains were transformed with one or two plasmids.

| Strain | Designation | Plasmids | Input | Coupling | Output |
|---|---|---|---|---|---|
| Cloning Strain | XL1-Blue | - | - | - | - |
| $I^-C^-O^-$ | EW11 | - | - | - | - |
| $I^-C^-O^+$ | EW11-JA01 | pSS9-RNA | - | - | CymA-MtrCAB |
| $I^{C42A}C^-O^-$ | EW11 | pFd007_C42A/lacI/fnr/pgl + p(e-)nzymes_NC | Fd(C42A) | - | - |
| $I^{C42A}C^-O^+$ | EW11-JA01 | pFd007_C42A/lacI/fnr/pgl + p(e-)nzymes_NC | Fd(C42A) | - | CymA-MtrCAB |
| SQR Analysis | EW11 | pFd007_C42A/lacI/fnr/pgl + p(e-)nzymes_NC | Fd(C42A) | - | - |
| SQR Analysis | EW11 | pFd007_C42A/lacI/fnr/pgl + p(e-)nzymes | Fd(C42A) | gsSQR | - |
| SQR Analysis | EW11 | pFd007_C42A/lacI/fnr/pgl + p(e-)nzymes_2 | Fd(C42A) | rcSQR | - |
| $I^{C42A}C^+O^+$ | EW11-JA01 | pFd007_C42A/lacI/fnr/pgl + p(e-)nzymes_2 | Fd(C42A) | rcSQR | CymA-MtrCAB |
| $I^+C^+O^+$ | EW11-JA01 | pFd007/lacI/fnr/pgl + p(e-)nzymes_2 | Fd | rcSQR | CymA-MtrCAB |
| $I^SC^+O^+$ | EW11-JA01 | pERA007.55/lacI/fnr/pgl + p(e-)nzymes_2 | sFd-55-ER | rcSQR | CymA-MtrCAB |
